# Agroecosystemic Resilience Index (AgRI): a method to assess agrobiodiversity

**DOI:** 10.1101/2020.12.03.409656

**Authors:** José-Alejandro Cleves-Leguízamo, Eva Youkhana, Javier Toro-Calderon

## Abstract

Agricultural production systems, subjects of study in agroecology, are non-equilibrium open systems permanently influenced by the action of natural or anthropogenic disturbances or “ripple effects.” Faced with this situation, agroecosystems tend to maintain a state of functional equilibrium in time and space, through an emergent property known as *resilience*. This concept is related to the dissipative capacity of agroecosystems to interact with the disturbance in such a way as to allow it to preserve its functionality and basic structure, through attenuation of the effect that disturbs the system. The literature reports diverse methods with a variable number of indicators or criteria for the evaluation or analysis of resilience. Many of these present conceptual deficiencies considering that the components of the system have similar characteristics and linear responses, that is, they do not show changes due to the action or nature of the disturbance. In this sense, there is a need to propose a generic method to analyze and evaluate agroecosystemic resilience, through a complex and comprehensive approach that takes into consideration the interaction of physical, biotic, socioeconomic or symbolic components of the system. These interactions are differential (weighted), to facilitate decision-making by the community, farmers, or administrators, regarding adaptations, adjustments or modifications that allow the agroecosystem to maintain its productivity and permanence.

## Introduction

Agroecosystems permanently interact with disturbances of natural and anthropic origin, i.e. the occurrence of adverse weather events, public policies, market fluctuations, community organizations, the role of institutions, availability of information and economic, logistical, technical, administrative or infrastructure resources, among others.

Faced with the occurrence of these events, the affected system responds to the disturbance dynamically, thanks to an emerging property of these open systems called resilience (1). In recent decades, this concept has motivated the development of numerous scientific investigations (2–10), due to multiple challenges experienced by sociological systems (11) and reflected in political turmoil, power relations, terrorism, famines, forced displacement, refugee crises, and high recurrence and intensity of extreme weather events. The effects of these are perceived among the most vulnerable populations, generating serious health effects, and changes in productivity, which have a fundamental impact on food security and sovereignty (12–14). Faced with this problem, scholars propose studying resilience under the systematic dynamics (SD) approach (15).

The concept of resilience is considered a property or a multidimensional and multidisciplinary phenomenon, with definitions and interpretations that vary according to the context, purpose, scale, system, discipline and domain (10). It was originally analyzed by engineers to analyze the ability of some materials to respond to the action of external forces or stress. In the 1950s, this term was initially introduced in psychology, defined as the ability of human beings to react, adapt and learn from critical situations (16).

Resilience has roots in socioecological theory from the concepts proposed by Holling (17), who indicates that from the environmental point of view, resilience provides a link between the theory and practice of ecology. Its evaluation and analysis (18) require a systemic and complex approach. For this reason, the formation of multidisciplinary work teams is recommended, to avoid partial and low-range solutions.

Resilience is a complex concept and a function of ecosystem attributes (with thermodynamic limits) and cultural attributes (without thermodynamic limits), the confluence of which configures the resilience of the socio-ecological systems (SES) of communities at the local or regional level (19). In the social context, resilience is related to the capacity for response, transformation, adaptation and innovation that socio-ecological systems (SES) have in the face of different natural and anthropogenic disturbances, in such a way that they adjust their structure to continue their productive function (20).

Conceptually, two types of resilience are considered. The first type, natural resilience, is more general and is associated with the genome and environmental supply. The second, acquired resilience, is more specific and closely linked to the nature of the disturbance (21). Regarding the analysis of the resilience of agroecosystems (12), the authors indicate that there is evidence of a direct relationship between biodiversity and resilience. Analysis and interpretation would allow the design and implementation of adaptations in agricultural systems, guaranteeing the production and availability of food in a timely manner according to populations’ needs.

From the point of view of methodologies and methods for evaluating agroecosystem resilience, the literature reports different approaches applicable in global territorial contexts, i.e. the Resilience Index Management and Analysis – (RIMA II) (22); the integrated Household Drought Risk Index - (HDRI) (23); the Index of Holistic Risk - (IHR) (24); the REDAGRES methodology (25); the Evaluation of Agroecological and Conventional System Resilience (26); the modified REDAGRES methodology (27,28); the methodology for the diagnostic of biodiversity complexity (29), the Index of Natural Resource Sensitivity to Drought (30); the Agroecosystems Performance Index - (API) (31); the Response Inducing Sustainability Evaluation method - (RISE) (32); the MESMIS method (33); and the Sustainability Assessment of Farming and the Environment Framework method - (SAFE) (34).

When analyzing these methods, restrictions are observed in their applicability. For example, for some, a high number of criteria or indicators are either duplicates or are too difficult to interpret. In other cases, key criteria related to biophysical, socioeconomic, and cultural components are absent. Most importantly, some methods report the same response among system components both before and after a disturbance. These limitations increase uncertainty regarding an analysis of agrobiodiversity resilience, creating difficulties in making decisions about the design and implementation of management measures that prevent or mitigate the effects of natural or anthropogenic disturbances.

With this in mind, the present article aims to propose a method for the analysis of agroecosystemic resilience, called Agroecosystemic Resilience Index (IRAg), developed by (35). The concept of method, which is handled in this article, is circumscribed to the approach of (36,37), who define methodology as the approach that shapes the body of knowledge of a certain subject of study, in this case the evaluation of agroecosystem resilience, and method, as the technical procedures applied for a given analysis.

The document is divided into five chapters: **the first** addresses the conceptual definition of resilience, including socio-ecological systems with an agroecological approach; **the second** describes the behavior of agroecosystems in the face of climate variability; **the third** presents an overview of the main methodologies for assessing resilience; **the fourth** presents the methodological development of the Agroecosystemic Resilience Index (AgRI); and **the fifth** demonstrates an application of the AgRI as well as the conclusions.

### Resilience

Resilience is considered a multidimensional emergent phenomenon or property whose analysis requires a multidisciplinary approach. The definition and interpretation of resilience varies by context, purpose, scale, system, discipline, and domain (10). Originally, this concept was proposed in the context of physics to describe the elasticity and plasticity of materials. In psychology studies, the concept was used to explain the ability to interact and adapt to adversity, trauma, or tragedy (16). (17) pioneered the application of this concept to ecological stability and defined it as “a measure of the persistence of systems and their capacity to absorb changes and disturbances and still maintain the same relationships between populations or variables of state.” This definition diverges from the term stability, which “represents the capacity of a system to return to a state of equilibrium after a temporary disturbance; the faster it returns and the less it fluctuates, the more stable it will be.”

In biology, the concept arose when environmental groups constructed hypotheses to explain why, when faced with a disturbance, some natural systems collapsed, and others did not (38). (19) defined resilience as a measure of the ability to absorb changes, presenting cyclical (slow and fast) and crossed movements called pentarchy, identifying adaptation as the key mechanism of resilience (39–41), and recognizing the importance of uncertainty in previous processes (42–44).

Resilience is also applicable to social systems, where it is defined as the ability of groups or communities to interact with external disturbances because of social, political, and biophysical changes (41,45–47). (46) raises two approaches to the concept: i) social resilience, defined as the capacity of individuals and communities to interact with physical, biotic, social and political disturbances, and ii) ecological resilience, defined as a characteristic of ecosystems that remain functional in the face of disturbances. It is important to note, for rural studies applying a territorial approach, that interactions of great relevance are generated between these two types of resilience.

(48) highlight that resilience refers to the limits of disturbance that a system can absorb before changing to an alternative state. In this process there are multiple equilibria, which lead to the structural reorganization of the system. This attribute is considered an emergent adaptive property that allows complex systems to regenerate and / or transform.

(49–53) argue that resilience is an emergent property, generated in complex systems. It is the result of the interaction of its components at different scales, which allows them to cushion, adapt and especially innovate and transform not only in the face of specific stress factors, but also in the face of the inevitable and continuous biophysical and social changes in the environment (35).

(17,40,41,54–56) define resilience as the characteristic of a system that allows it to absorb disturbances, maintaining its structure and function, as well as ecosystem services. Regarding resilience and climatic events, (57) argues that resilience is related to the ecological niche of the species, the extent of the climatic anomaly, the association with mesoclimatic phenomena, exposure, vulnerability and the capacity for adaptation.

For (53), resilience is associated with three specific system capacities:

a. adaptation, defined by (41) as the capacity of the system to respond to external changes, while maintaining its operation. At the community level, it requires ingenuity, the ability to identify problems, set priorities, mobilize resources, and combine experiences and knowledge to adjust responses in a changing context (35,58).
b. damping, a response of the system that allows it to dissipate the energy caused by a disturbance, allowing it to preserve its integrity and functionality (59). This characteristic of the systems is identified by (60) as a dissipative structure.
c. transformation, understood as the capacity of a system to change to new structures and operations, when the previous system is not sustainable (61).

(62) argue that vulnerability and resilience are inversely proportional. In this sense, the vulnerability of a system increases as its resilience decreases (Fig. 1).

**Figure 1.**
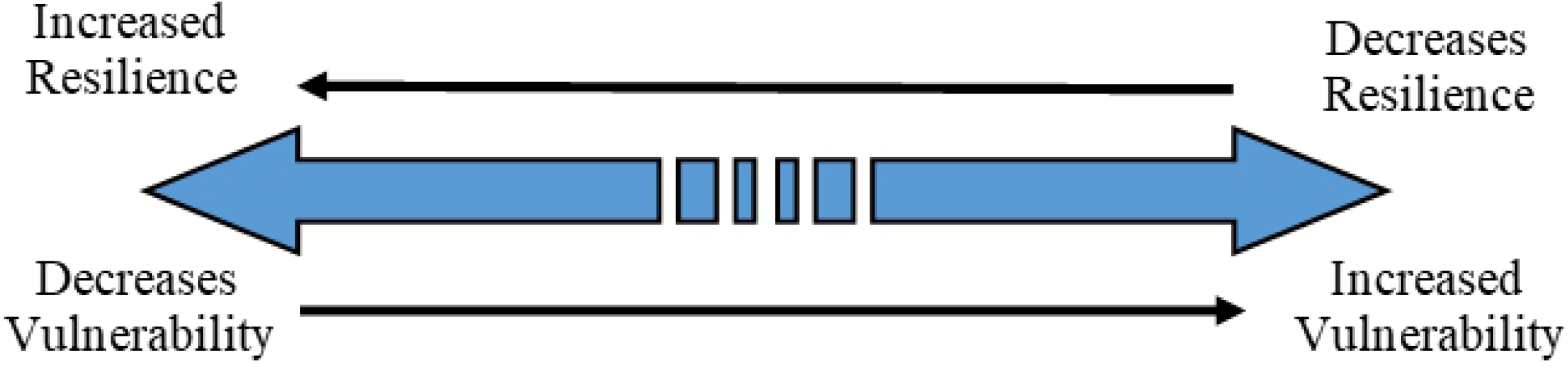
Relationship between vulnerability and resilience (modified from (62))

There is a great number of definitions. However, they converge in that resilience is an emergent property, which allows systems to adapt in response to the influence of disturbances. This characteristic is the object of interdisciplinary research due to the physical, biotic, social, economic, and cultural repercussions of natural and anthropogenic phenomena that threaten the life and well-being of individuals, communities and ecosystems, which are increasingly recurrent and intense (10).

### Resilience of Social-Ecological Systems (SES) and agroecosystems

The analysis of the relationships between the social and ecological components in a territory gives rise to the formation of Social-Ecological Systems (SES). This concept arises from systematic studies with multi and interdisciplinary approaches and has become the analytical framework for studying interactions within anthropic natural systems. Thanks to work developed by (20,49,63), SES is being widely used in various knowledge areas such as human and natural sciences, engineering, arts and medicine (64,65), although it is a relatively new concept. (64) found 2,165 publications related to SES between the years 1998-2016.

The concept of SES was initially proposed by (66) in his Doctoral Thesis “Applications of engineering systems. Analysis to the human social-ecological system.” Later, (67) used this concept in a publication on epidemiology. However, it was (63) who, in their work on resilience, proposed a framework for the study of ecosystems and human institutions called socio-ecological resilience, in the context of The Millennium Ecosystem Assessment. (68) used this concept in the framework of interdisciplinary studies (64,65). In 2017, (69) contributed to the evolution of the SES theoretical framework, recommending the use of the term social-ecological system, instead of socio-ecological system, arguing the following: “social-ecological emphasizes that the two subsystems are equally important, whereas socio- is a modifier, implying a less than equal status of the social subsystem” (69). Figure 2 presents a timeline of the main contributions to the construction of the SES theoretical framework.

**Figure 2.**
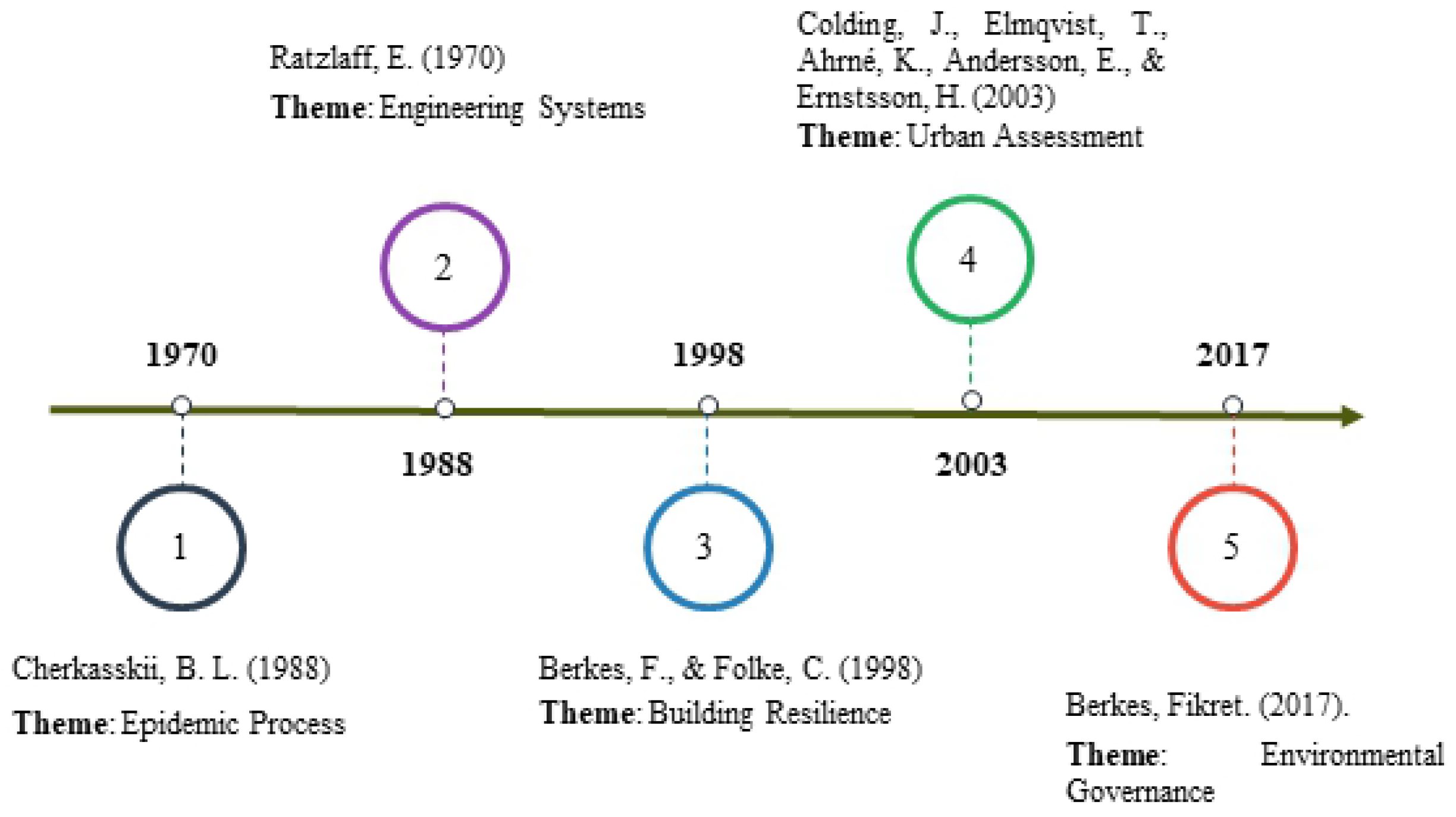
Timeline of the construction of the SES theoretical framework (64,65)

Regarding social-ecological systems, there is a great number of complementary concepts. However, (70) state a series of peculiarities that allow for their characterization. They argue that SES are complex adaptive systems with the following characteristics: “i) integrated biogeophysical and socio-cultural processes, ii) self-organization, iii) nonlinear and unpredictable dynamics, iv) feedback between social and ecological processes, v) changing behavior in space (spatial thresholds) and time (time thresholds), vi) legacy behavioral effects with outcomes at very different time scales, vii) emergent properties, and viii) the impossibility to extrapolate information from one SES to another” (70).

In agroecosystem analysis, (28) propose a complementary concept by stating that resilience is an emergent property that is not intrinsically neutral, positive or negative “due to the lack of consensus in society on the objectives and strategies to respond or interact with changes or disturbances”. Likewise, this concept has been associated with power, increasing in social groups with greater access to resources and political participation, and decreasing in groups with less economic capacity (71).

(72–75) mention that the reduction of social vulnerability through the transfer of knowledge and technology, the promotion of cultural processes of collective self-organization, as well as the adjustment of state institutions, contributes to increasing agroecosystem resilience.

By virtue of the structural and functional organization level, agroecosystems will be more or less responsive to a disturbance. In this way, the level of information is definitive, as well as farmers’ resource availability.

(76) propose incorporating resilience within the framework of rural development, to radically transform it in favor of environmental justice. Resilience can support communities’ rights to food sovereignty, specifically in populations vulnerable to climate change, introducing the concept of people-centered resilience (social resilience for development).

According to (62), resilience can have two categories: i) innate or own resilience that depends on the natural characteristics of the system, and ii) acquired resilience, built from physical, biotic, social, economic and cultural adaptations.

In this document, an agroecological approach to resilience is defined as an emerging attribute of agricultural systems, which allows them to interact, respond and adapt to disturbances of natural or anthropic origin. These systems react through autopoiesis, the democratization of available resources, technical assistance, market articulation, and available infrastructure. This includes the incorporation of ancestral knowledge and practices that promote agrobiodiversity through an articulation with the principal ecological landscape structure, the management of vegetation cover and water and soil conservation processes. This is done in a way that allows farmers and communities to continue with their traditional production processes, based on their culture, food autonomy and well-being (28,35) (Fig. 3).

**Figure 3.**
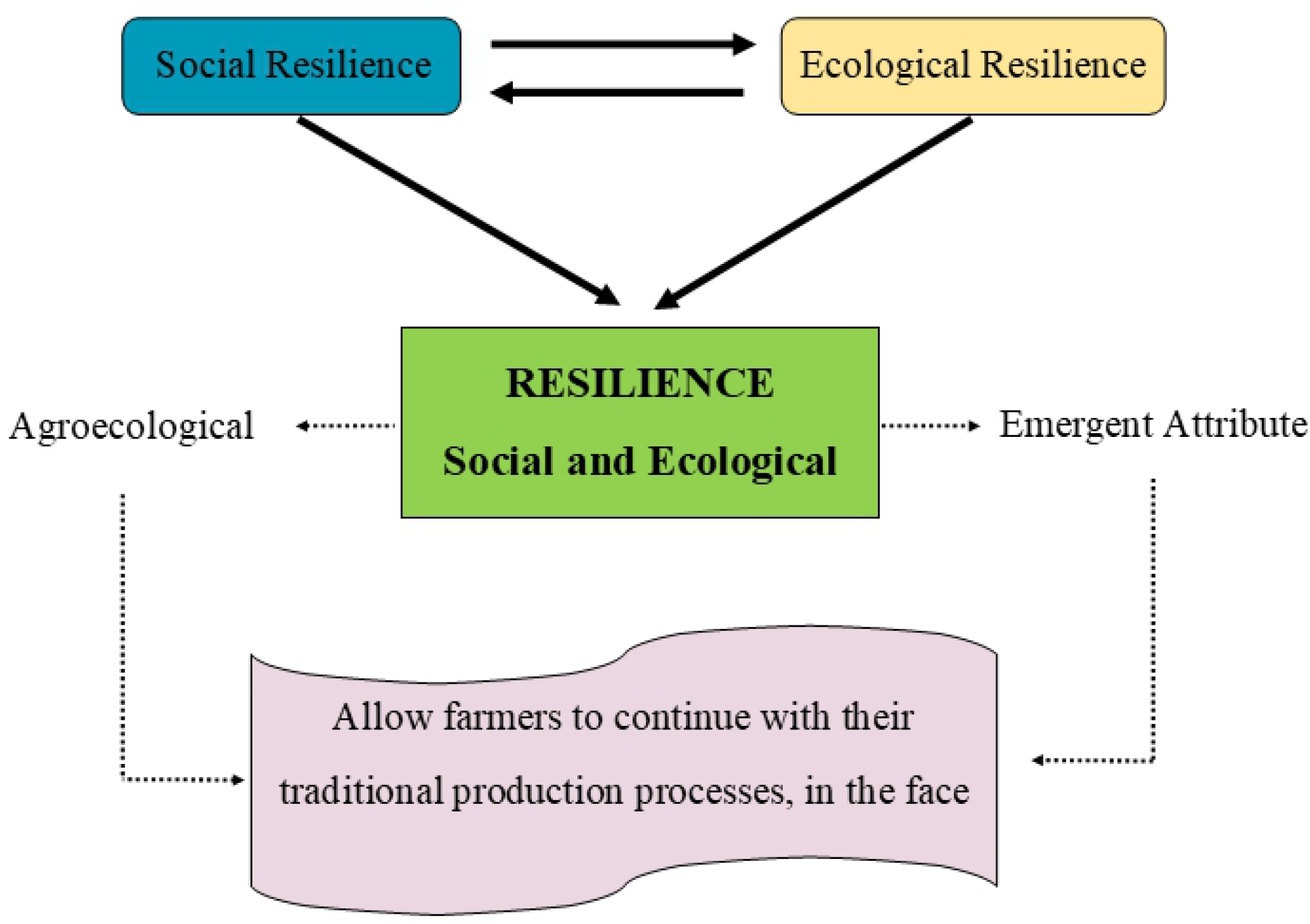
The concept of resilience with an agroecological focus.

## METHODS FOR EVALUATING RESILIENCE

The assessment of resilience must include the complexity of the ecosystem and cultural interrelationships that define agroecosystems. This work, at the Latin American level, has recently been approached through Indices that try to capture the greatest number of variables.

However, these methods present limitations from the conceptual and methodological point of view, especially related to the non-incorporation of elements of the social system and the perception that the elements of the agroecosystem interact under equal conditions before and after disturbances. The main methods are described below.

### The FAO’s Resilience Index Measurement and Analysis Model (RIMA-II)

To evaluate resilience, the Food and Agriculture Organization (FAO) has proposed and implemented models with qualitative and quantitative approaches. One of the most representative is the FAO’s Resilience Index Measurement and Analysis (RIMA-II). This index conceives resilience as the ability of a household to recover from a disturbance and is linked to food security and sovereignty (22). This index has been used in several African countries including Ethiopia, Kenya, Sudan, and Somalia.

The FAO considers that resilience can be evaluated or analyzed in two ways: i) direct: that is, in a descriptive (generic) way, which is useful for proposing public policies, and ii) indirect: using the variable model MIMIC (Multiple Indicators, Multiple Causes) based on statistical techniques of multiple comparison that allow observations to be made over time.

Researchers have combined the direct and indirect measurements by proposing the Resilience Capacity Index (RCI) (22).

In this context, the assessment of resilience helps to understand and support the capacity of households (the basic unit of the food system), to recover from natural and social events.

The index proposed by FAO is based on two dimensions: i) the physical dimension and ii) the response capacity, grouped into an equation with nine unweighted indicators which are subsequently modeled and analyzed.

### The Integrated Household Drought Risk Index (HDRI)

This method was proposed by Luetkemeier & Liehr, (2018) in sub-Saharan Africa, specifically in the municipality of Cuvelai, located in the river Cunene basin (Angola and Namibia), to evaluate the vulnerability of households in different socioeconomic and environmental contexts. It formulates a composite indicator with three general dimensions (risk, sensitivity, and response capacity) and 13 indicators for variable analysis, including precipitation supply, water demand, edaphologic characteristics, and the institutional supply of infrastructure and financing.

### Index of Holistic Risk (IHR)

This method is an adaptation of the proposal of (24), for social research within the framework of natural disasters of climatic origin.

This index was evaluated in Latin America by (77), analyzing the threat of environmental risks generated by climate change (especially the recurrence and intensity of droughts). For this purpose, ten variables were identified in relation to indicators of threat, vulnerability and response capacity in peasant and indigenous productive systems in the region of the Chilean Araucanía.

The main limitation of IHR is that it does not identify the variables with greatest specific relevance since it does not perform a weighting of variables.

### REDAGRES Method

The REDAGRES method was proposed by (25) and applied in several countries in South America (Brazil, Chile, Colombia, Cuba, Mexico and Peru). It estimates non-weighted ecosystem factors and socio-cultural variables (social organization, solidarity networks, traditional knowledge) that enhance, limit, or explain the resilience of agricultural systems.

It groups 55 criteria into the following categories: physical (four), soil (five), water management (eight), biological diversity (nine), social aspects (thirteen), economic aspects (seven), institutional aspects (six), and technological aspects (three). The main objective of this method is to estimate resilience against the occurrence of catastrophic climatic events such as storms, hurricanes, and droughts.

The direct summation is represented on a scale of 1, 3, and 5, where 1 indicates low, 3 intermediate, and 5 high resilience of socio-ecological systems.

The results are presented in the form of a traffic light for the prioritization of future activities or management practices, where green is optimal, yellow is medium and red is low resilience (35).

### Evaluation of agroecological and conventional system resilience

This method was developed under the approach of community participation by a team comprised of technicians and representatives of the peasant community within the social and ecological context of Colombia.

It was applied by (26) based on the methodology proposed by (25) who compare cultural management practices with an agroecological and conventional approach within the department of Antioquia (Colombia), using a mixture of empirical methods and rapid agroecological evaluation.

The selected indicators approached the threat of the occurrence of adverse climatic events and evaluated the level of vulnerability based on the Wilcoxon test (78), estimating the capacity for recuperation or the cultural response of farmers in their parcels through the introduction of adjustments in practices of management techniques in their crops.

The method evaluates six physical indicators of vulnerability: i) slope, ii) landscape diversity, iii) infiltration capacity, iv) analysis of the biostructure, v) gullies and vi) level of soil compaction and thirteen indicators of response capacity: i) coverage, ii) living fences, iii) conservation tillage, iv) water management, v) organic matter management, vi) contour lines, vii) food for self-consumption, viii) use of inputs, ix) seed bank, x) animal feed, xi) crop association, xii) protection areas and xiii) soil textural class. All estimates are calculated without weighting.

With the response capacity and vulnerability index, a risk index (RI) was generated, allowing to establish that local adaptive management (cultural) practices are a key strategy for the construction of socio-ecological resilience. The calculation of risk is presented in (Eq. 1).

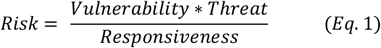

According to the vulnerability evaluation, every indicator is assigned a value based on a “traffic light” approach to propose corrective actions. The numeric values were assigned within a range of 1-5 in the following manner: 1 (green), 3 (yellow) and 5 (red).

### Modified REDAGRES method

The modified method was proposed by (27) to evaluate resilience based on environmental characteristics (ecosystemic and cultural), as well as farmers’ management practices. The method aims to measure resistance and recoverery from the effects caused by the incidence of climatic variability in agroecosystems, mainly of conventional and agroecological coffee growers in the Colombian context.

The method analyzes 10 categories grouped into 64 criteria, organized into 4 physical parameters; 5 edaphic characteristics; 8 water management practices; 9 paramentos of biological diversity; 13 social aspects; 7 economic aspects; 6 institutional aspects; 3 political aspects and 3 technological levels (35).

This method introduces adjustments to the REDAGRES method by including new parameters in the cultural, social, and economic traits, as well as in the management and conservation practices of water and soils.

Similarly, the variables are given the same importance, that is, there are no weights. The results are presented in a traffic light format, proposing future management measures.

### Method to evaluate the Principal Agroecological Structure (PAS)

This method was proposed at the National University of Colombia by León-Sicard in the first decade of the year 2000 (79,80). The PAS is an indicator of the agrobiodiversity of the major (farm) and minor (lot) agroecosystems.

In this context, agroecosystems respond differently to disturbances of anthropogenic and natural origin. This interaction is related to the physical, chemical, and microbiological properties of the soils, the area and slope of the farm, the presence of restrictive geological components as well as other ecosystem characteristics. Sociocultural variables such as land tenure, farm size, educational level, family structure, knowledge about climate behavior and production systems are equally fundamental.

The resilience of agroecosystems is directly related to agrobiodiversity. According to (81) the design and productive arrangement of the most diverse agroecosystems generates greater resilience. Among the practices successfully implemented in these agroecosystems, we can mention silvopastoral systems (SSP), intercropping, associated crops and, in general, polycultures.

Taking into account this conceptual approach, the PAS analysis method assesses five ecosystem attributes without weights: i) Connection with the Principal Ecological Landscape Structure (CPELS); ii) Extension of External Connectors (EEC); iii) Diversification of External Connectors (DEC); iv) Extension of internal connectors (EIC); v) Diversification of Internal Connectors (DIC) and five cultural attributes: i) Land use (LU); ii) Weed Management (WM); iii) Other Management Practices (OP); iv) Perception - consciousness (PC) and v) Capacity for action (CA).

The 10 attributes are assigned a rating scale ranging from 1 to 10 in such a way that the highest rating is 100.

The results obtained have the following interpretive scale: 80-100 (Strongly developed); 60-80 (Moderately developed); 40-60 (Slightly developed); 20-40 (weakly developed) and <20 (without structure).

Studies have shown that agroecosystems with strongly developed PAS show greater resilience against the occurrence of disturbances of diverse natures (79).

### Method to estimate the Biodiversity Management Coefficient (BMC)

The BMC evaluates the resilience to climate change, analyzing the complexity of the designs and cultural management of biodiversity.

It proposes 64 indicators, grouped into the following categories: 18 indicators of productive biodiversity and management of the elements of productive biodiversity (MPrB); 7 on soil management and conservation (SMC); 5 related to water management and conservation (WMC); 5 indicators for phytosanitary intervention management in productive areas (PIMPr); 15 on the design and management of the elements of auxiliary biodiversity (DMAuB) and 14 on the status of the elements of associated biodiversity (EABs).

Similarly to the PAS, the results obtained allow an estimation of the Biodiversity Management Coefficient (BMC), an indirect measure of resilience, based on the following scale: 0.1-1.0 Simplified; 1.1-2.0 Low complexity; 2.1-3.0 Medium complexity; 3.1-3.5 Complex; 3.6-4.0 Highly complex (29,35).

### Method to evaluate the Index of Sensitivity of Natural Resources to drought (SNRs)

In order to analyze the resilience of production systems in the face of drought from an agroecological perspective, (30) proposed a conceptual and methodological framework to evaluate unweighted agroecological indicators, based on resistance-absorption functions (AR), recovery (RC) and transformability (TR).

The Index of Sensitivity of Natural Resources (SRNs) was constructed through the analyzed variables. This index focuses on scales based on farm-level sensitivity of crops, animals, soils, as well as water availability under conditions of water stress. The equation to evaluate the General Drought Resilience Index (DRIs) of each productive unit is presented in (Eq. 2):

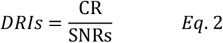

Where: CR= Capacity of resilience of farm-level systems; SNRs= Sensitivity of natural resources to drought.

### Method to assess the disaster resilience index (CDRI)

It is important to assess resilience in the event of natural disasters to enable proper irrigation management. To this end, scholars proposed the CDRI method, which includes 6 categories: i) access to services and quality of institutions, ii) quality of housing, iii) social cohesion, iv) educational level, v) environment and vi) availability of resources. Within these categories there are 8 subcategories, where 28 indicators are grouped. The most suitable technique for normalization is the standardization method (AMP) proposed by (82).

In Italy, the index was useful for developing quantitative assessments and identifying patterns of social vulnerability and differences in resilience between northern and southern regions, to propose adaptation measures for climate change (83).

When (84) analyzed the results of 174 publications aimed at evaluating resilience to natural disasters between 2005 and 2017, they found that 39.7% of the articles used qualitative methods, 39.1% quantitative methods and only 10.3% performed practical validation of the proposed resilience indices.

### Agro-ecosystem Performance Index (API)

The API index was proposed as a comprehensive method, developed with community participation, to evaluate the performance of agroecosystems in terms of sustainability.

The index integrates quantitative sustainability indicators, through two socio-economic and one ecosystem category: i) economic aspects, ii) social aspects, and iii) conservation of natural resources. Finally, a statistical analysis of principal components (PCA) is carried out.

This index was used to assess the sustainability of rice production systems in Kerala, India and to propose adjustment measures, to establish development priorities for local communities (31).

### RISE (Response, Induction, Sustainability, Evaluation) Method

The RISE method was proposed in order to analyze the sustainability of the components of productive systems in small plots, so that farmers or communities immersed in a territory could recognize climatic effects and productive limitations, and propose corrective measures that allow them to increase resilience before the occurrence of natural disturbances.

The method evaluates 12 indicators grouped into 57 parameters: i) energy, ii) water, iii) soil, iv) biodiversity v) emissions, vi) crops, vii) waste, viii) flows, xix) income, x) investment, xi) economy and xii) local situation. The method quantifies agroecosystems at the farm level in terms of their physical, biotic, economic and social dimensions (32,85,86).

### MESMIS Method (Framework for the Evaluation of Natural Resource Management Systems Incorporating Sustainability Indicators)

The MESMIS method is a useful analytical tool for the diagnosis of the state of production systems. It considers the local factor as a fundamental aspect of the diagnosis, offering endogenous responses. Under these conditions, it serves as a guide to organize and prioritize the activities to be implemented.

The method seeks to understand, in a comprehensive way, the limitations and possibilities for the sustainability of management systems that arise from the intersection of environmental, social, and economic processes. It analyzes numerous non-weighted indicators in a participatory way, among which we can mention productivity, stability, reliability, resilience, adaptability, equity, and self-management (33).

### SAFE (Sustainability Assessment of Farming and the Environment Framework) Method

This method evaluates agricultural activities through a hierarchical structure adapted from the application of the PCI theory (Principle-Criterion-Indicator). It analyzes economic, social and ecosystem indicators, considering the multifunctional character of agroecosystems to achieve sustainability (34).

As can be seen in each of the methods presented, the evaluation of resilience is a complex process, which intertwines aspects related to vulnerability, sensitivity, self-organization, and adaptation capacity. For this reason, operationalization and estimation proposals acquire great relevance with time (87). A summary of these methods is presented in Table 1.

When analyzing the methodologies, the following weaknesses or opportunities for improvement could be evidenced:

- High number of components and synthetic variables.
- Simultaneous evaluation of components in different categories.
- Lack of weighting of the components makes analysis and applicability difficult.
- Assuming that system components have identical, linear response capacities, ignores the reality that the components of any system have their own differential attributes associated with their own nature and composition.
- Lack of validation.

**Table 1.** Comparison of methods for assessing resilience

| Name of method | Author | Country/Region | Category | Indicators | Type of analysis | Use of Delphi methodology | Weight |
| --- | --- | --- | --- | --- | --- | --- | --- |
| Resilience Index Measurement and Analysis (RIMA- II) | (22) | Africa | 2 | 9 | Linear | NO | NO |
| Índice Integrado de Riesgo a la Sequía en los Hogares (HDRI). | (23) | África | 3 | 13 | Lineal | NO | NO |
| Holistic Risk Index (HRI) | (24) | South America | 2 | 10 | Linear | NO | NO |
| Resilience in agroecological and conventional systems Index | (26) | South America | 2 | 19 | Linear | NO | NO |
| REDAGRES method | (25) | South America | 8 | 55 | Linear | NO | NO |
| Modified REDAGRES method | (27) | South America | 10 | 64 | Linear | NO | NO |
| Method to evaluate the Principal Agroecological Structure (PAS) | (79) | South America | 2 | 10 | Linear | NO | NO |
| Biodiversity complexity Index | (29) | Central America | 6 | 64 | Linear | NO | NO |
| Sensitivity of Natural Resources to drought Index (SNRs) | (30) | Central America | 4 | 29 | Linear | NO | NO |
| Disaster resilience index (CDRI) | (83) | North America | 6 | 28 | Linear | NO |  |
| Agro-ecosystem Performance Index API | (31) | Asia | 2 | 3 | Linear | NO | NO |
| Response, Induction, Sustainability, Evaluation (RISE) | (32) | North America<br>Asia | 12 | 57 | Linear | NO | NO |
| MESMIS | (33) | South and North America | 7 | 30 | Linear | NO | NO |
| Sustainability Assessment of Farming and the environment Framework. (SAFE) | (34) | North America | 3 | 32 | Linear | NO | NO |

Considering these limitations, the Agroecosystem Resilience Index (AgRI) is proposed, the foundation and structure of which is described below.

## METHODOLOGY

Next, the methodology developed to obtain the Agroecosystemic Resilience Index (AgRI) is described. This index proposes a comprehensive evaluation of the ecosystemic and cultural components. The method is conceptually supported in ecosystem-culture or society-nature relationships and consider complexity, as well as decision-making in the context of environmental uncertainties (88).

The main methodological contribution is the use of qualitative and quantitative indicators generated from the weighted analysis of the main attributes of agricultural systems, recognizing that this approach improves prediction (89).

The method aims to identify the adjustments required to strengthen the components of the system with some degree of affectation, in such a way that their vulnerability can be reduced and the production and conservation of resources guaranteed.

### Methodological structure

Based on the structural methodological guidelines proposed by (90–95), the methodology for calculating AgRI is made up of the following phases: i) selection of categories, components and parameters; ii) weighting of the categories, components and parameters; iii) assignment of the interpretation scales of the parameters; iv) equation for the calculation of the Agroecosystemic Resilience Index (**AgRI**); and finally, v) interpretation of the Agroecosystemic Resilience Index (**AgRI**).

### Selection of categories, components, and parameters

A hierarchical structure of the environment was built based on the review of the methodologies presented in the previous chapter, conceptualized from the perspective of complex relationships between social, cultural, and natural components. This structure was divided into five categories made up of 13 components and 40 parameters. The five categories include the following: i) Ecophysiological, ii) Biotic, iii) Sociocultural, iv) Economic and v) Technological (Fig. 4).

**Figure 4.**
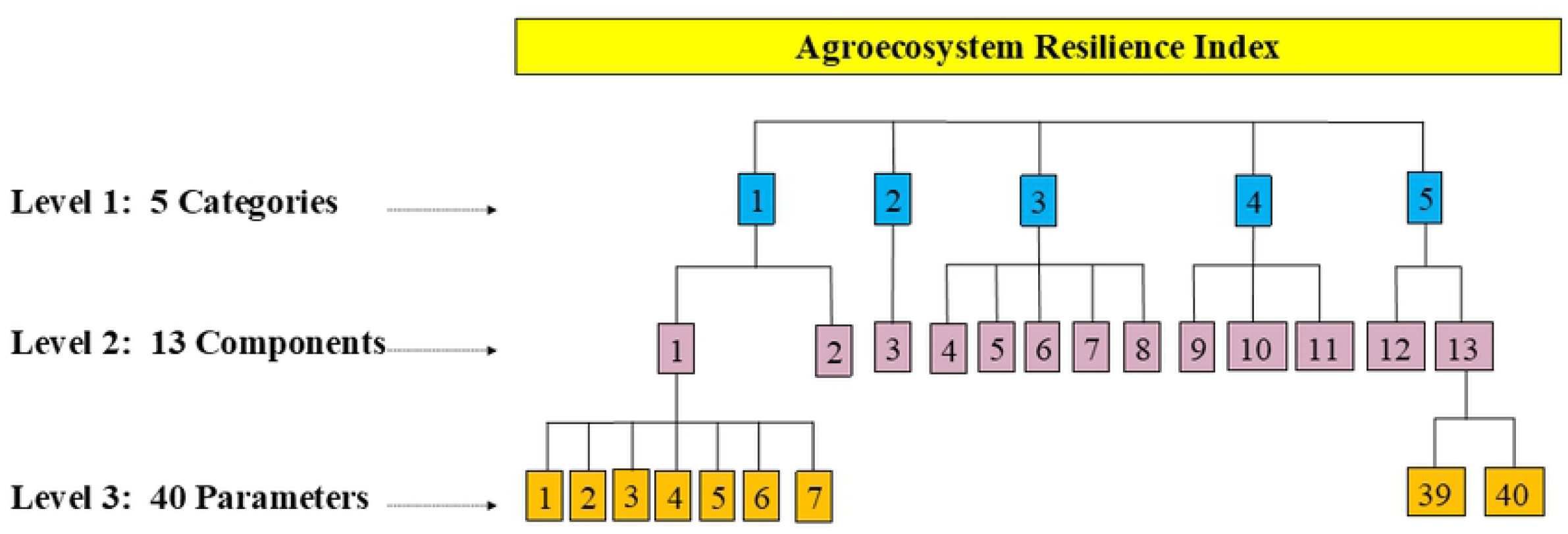
Hierarchical structure of AgRI.

### Weighting of categories, components, and parameters

As evidenced in the previous chapter, most of the methods for evaluating the resilience of agroecological systems do not take into account that the components of the environment interact with disturbances in diverse ways, according to innate or acquired resilience. In this sense, drawing attention to the fact that the proposed methods include a weighting process adjusted to this concept, (21,96) state: “the weighting of the indicators becomes a very important issue because not all indicators have to be equally important to explain the underlying sustainability phenomenon.” Therefore, higher weights are assigned to indicators that are more important than others, even though the weights can have a significant effect on the overall composite index.

Considering this recommendation and applying the Delphi method, proposed by (97) and widely used in scientific research (98), three rounds of consultation were carried out with 30 qualified experts. The experts held more than 15 years of work and academic experience in the areas of soils, crop protection, plant physiology, plant breeding, animal production, agricultural economics, rural sociology, zootechnics, edaphology, agrology and agricultural business administration. Consultations informed weights given to each of the environmental parameters (physical, biotic, social, economic, and cultural). Using the Google^®^ platform, a first round of surveys was sent, in which initial weighting values were proposed and modified for each of the reveiewed categories, according to their criteria.

During a second round, the previous results were sent for another round of modifications, to obtain the adjusted weights for the components.

During a third round, the results obtained through the proposed weighting of the categories, components and parameters were indicated. These were finally modified with an adjusted weighting (Table 2).

**Table 2.**
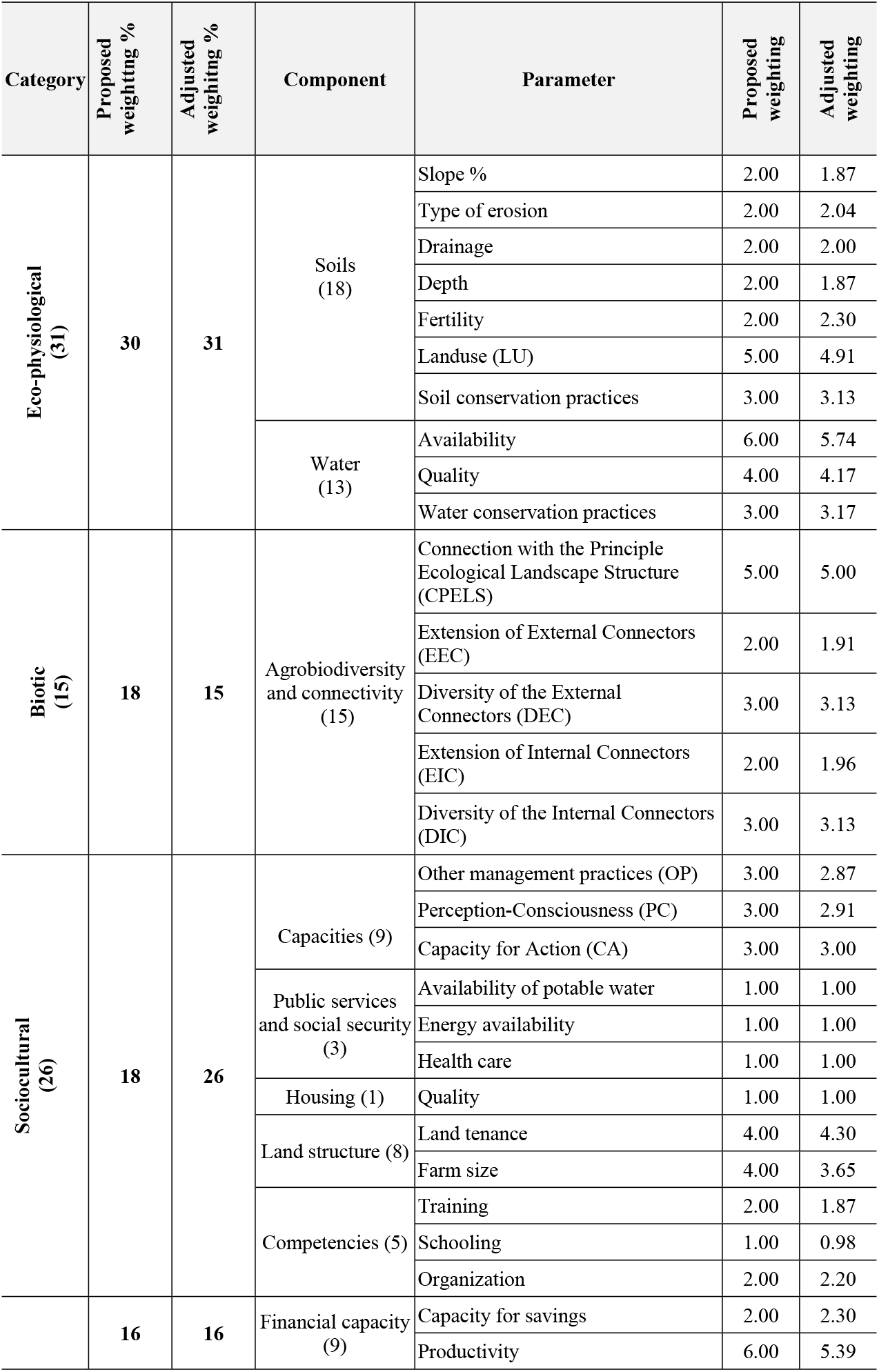
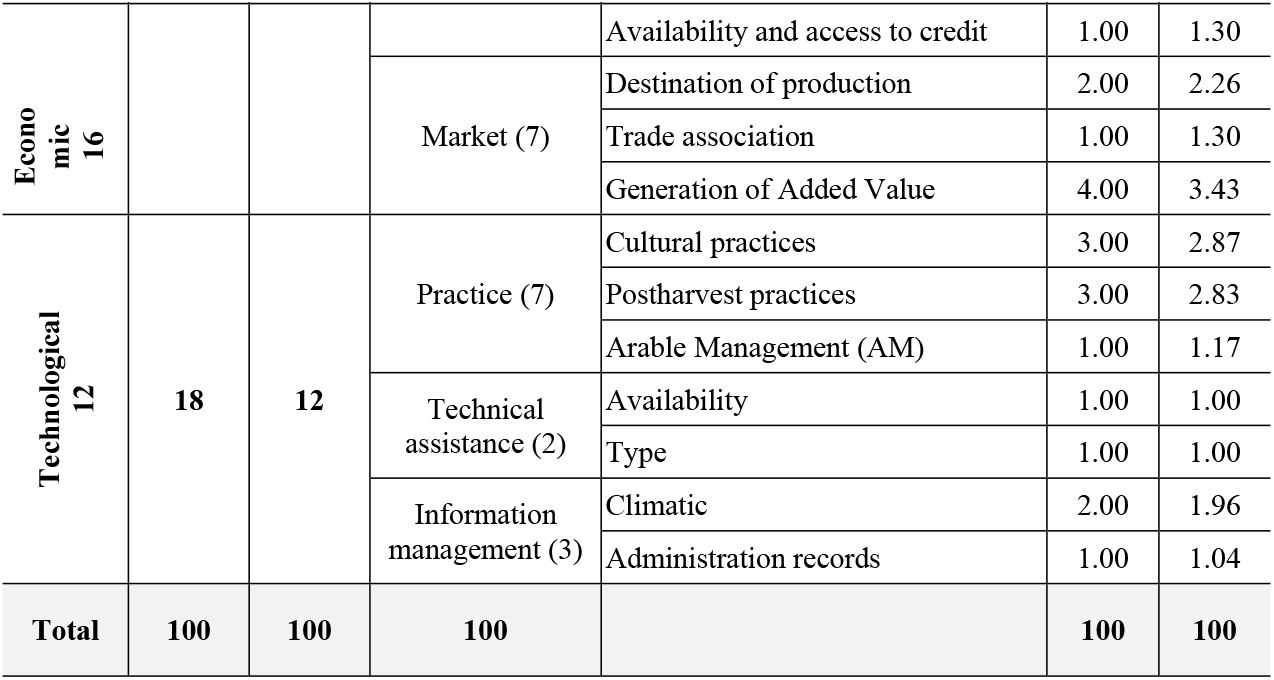
Weighting of categories, components and parameters (35).

It can be observed that the experts gave greater specific weight to the variables grouped in the ecophysiological category (31%), highlighting the availability of water resources (5.74) as well as water quality for irrigation (4.17). This is explained by the fundamental and irreplaceable role of water in all biotic processes. Soil use (4.91) stands out in terms of edaphic components, as a parameter that associates the cultivation types (mono or polyculture) with cultivation systems (spatial and temporal ordering). Agroecosystem design, as well as water availability, are definitive characteristics in the resilience capacity of agricultural systems.

The second category with the highest specific weight was the sociocultural category with 26 units, integrating the response capacity of farmers with service and infrastructure availability. The type of land tenure is highlighted (4.30) since a tenant does not have the same conservation interests as a landlord. Farm size also stands out (3.65). This expresses the difficulties that small farmers face when making intensive use of their farm area. Limits to crop rotation and conservation practices are relevant aspects in the capacity to respond to the effects of climate variability.

The economic category is associated with the availability of credit and the generation of economic surpluses that allow for the introduction of technological improvements, increased productivity (5.39) and the generation of added value (3.43).

As an expression of the biotic category, connectivity, and agrobiodiversity (PAS) demonstrates the importance of improving the level of connectivity of the minor agroecosystem with the surrounding landscape. Experts assigned 15% to this category.

For the technological category, the management of climate information at the farm level stands out (1.96). This is an aspect in which different institutions have been working with greater intensity in recent years, although with limited resources (35).

### Assignment of interpretive scales to the parameters

Taking into account the results obtained in the three rounds of the expert consultations, each parameter was assigned a rating, according to a qualitative interpretive scale in order to facilitate its interpretation, using the values of 1, 3 and 5. The rating of 5 is associated with attributes of high resilience, 3 with medium resilience and 1 with low resilience (Table 3).

**Table 3.**
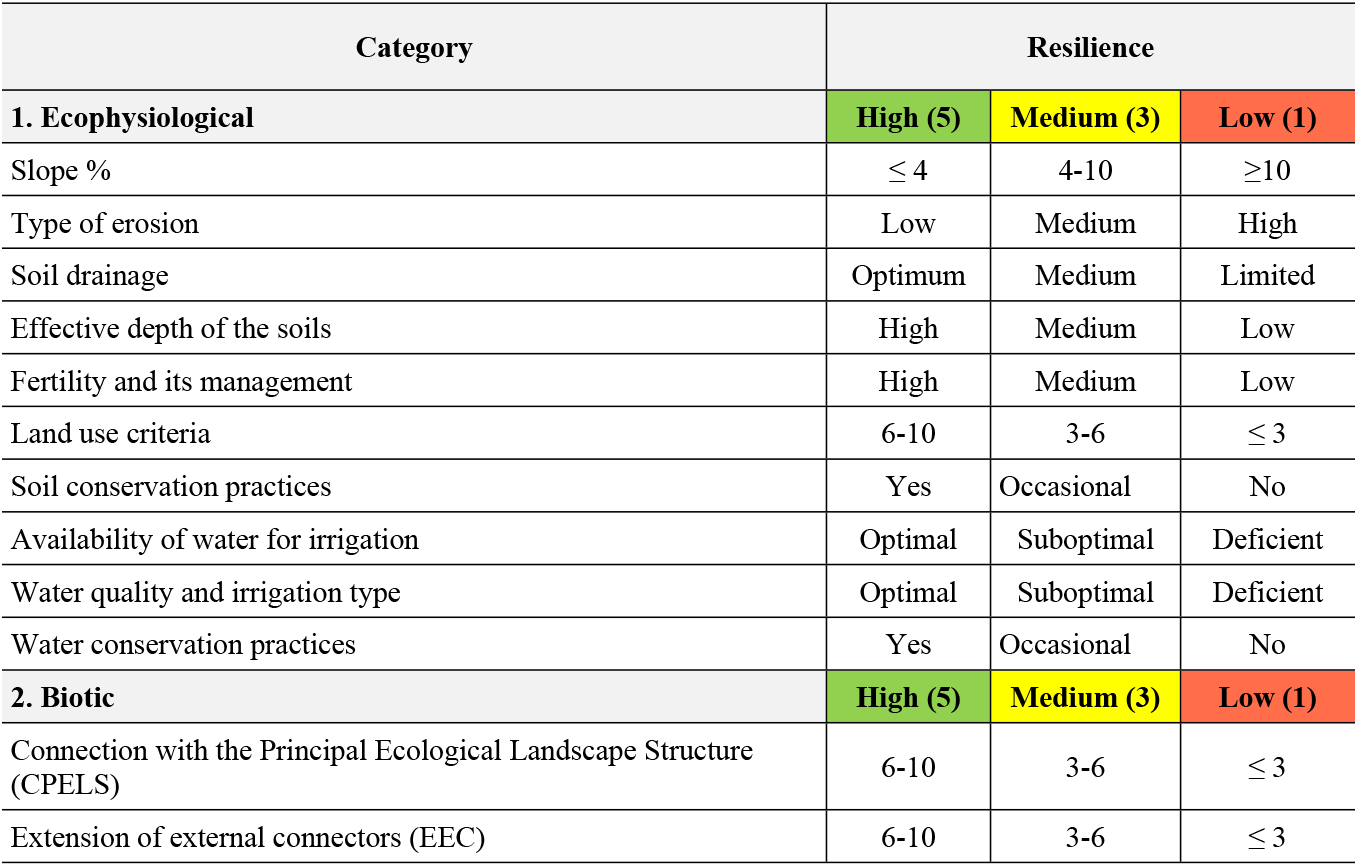
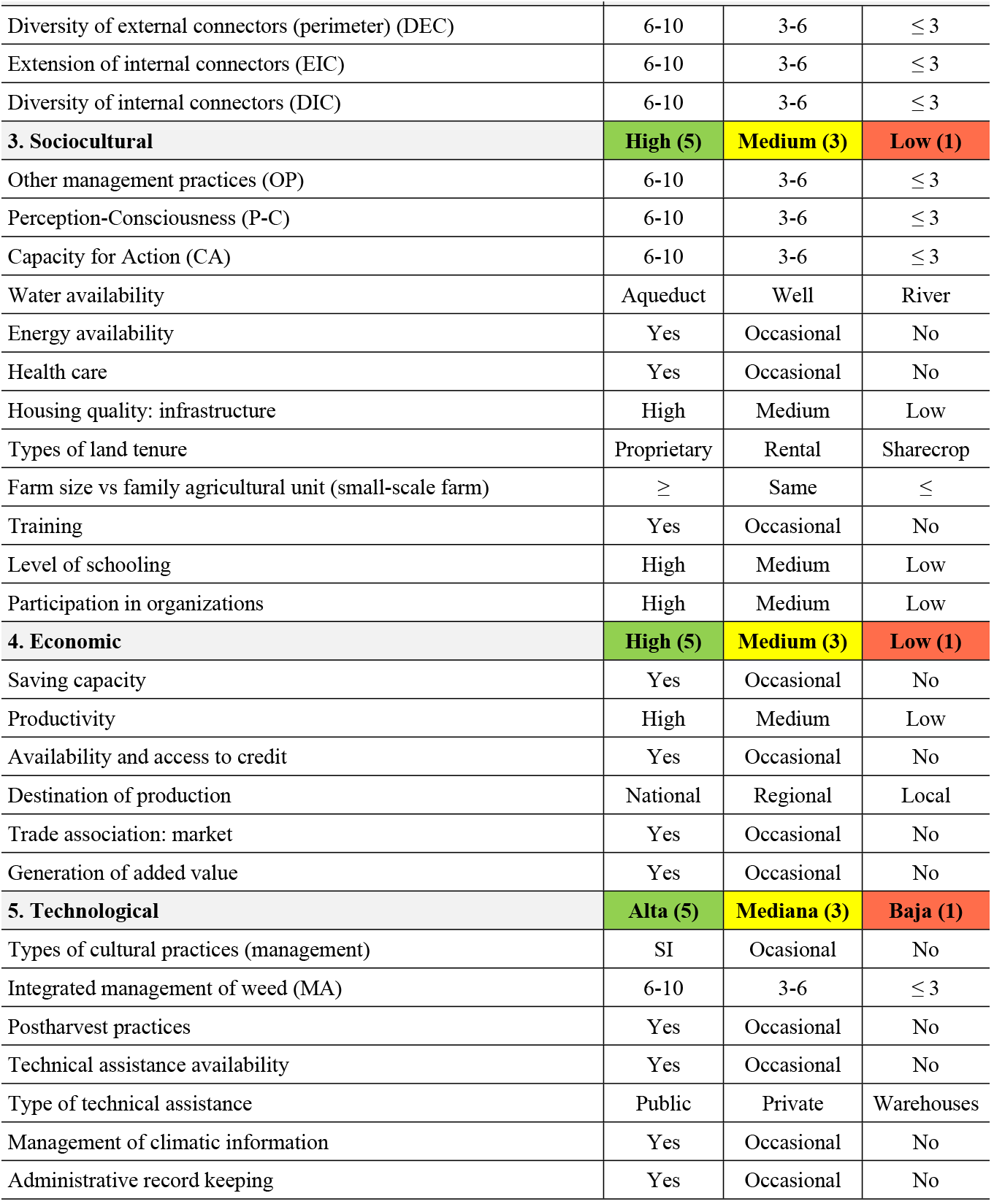
Interpretive scales for parameters (35).

### Equation for the calculation of the Agroecosystem Resilience Index (AgRI)

The Agroecosystem Resilience Index (ARI) is calculated from an equation (Eq. 1) whose terms correspond to the five (5) categories, which in turn group the forty (40) features that make up the agroecosystem (Fig. 4; Table 2).

Each parameter was weighted considering its potential resilience. The weights were obtained from the results obtained in two continuous rounds of expert consultations. The sum of the weighting of each group of parameters corresponds to the weighting of each category (Table 2), where the weightings of each of the categories, components and parameters are indicated in detail. The final equation for calculating the AgRI is presented in Equation 3.

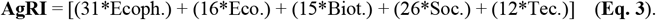

Where: **R** = Resilience; **Ecoph** = Ecophysiological Component; **Eco** = Ecosystemic Component; **Biot**. = Biotic Component; **Soc** = Sociocultural Component; **Tec** = Technological Component

### Interpretation of the Agroecosystemic Resilience Index (AgRI)

The AgRI corresponds to the weighted sum of 40 parameters, generating a result of 100 units. Each result is evaluated on a scale between 1, 3 and 5; therefore, the grade obtained will be in a range between 100 and 500, which is interpreted according to the parameters presented in **Table 4**.

**Table 4.**
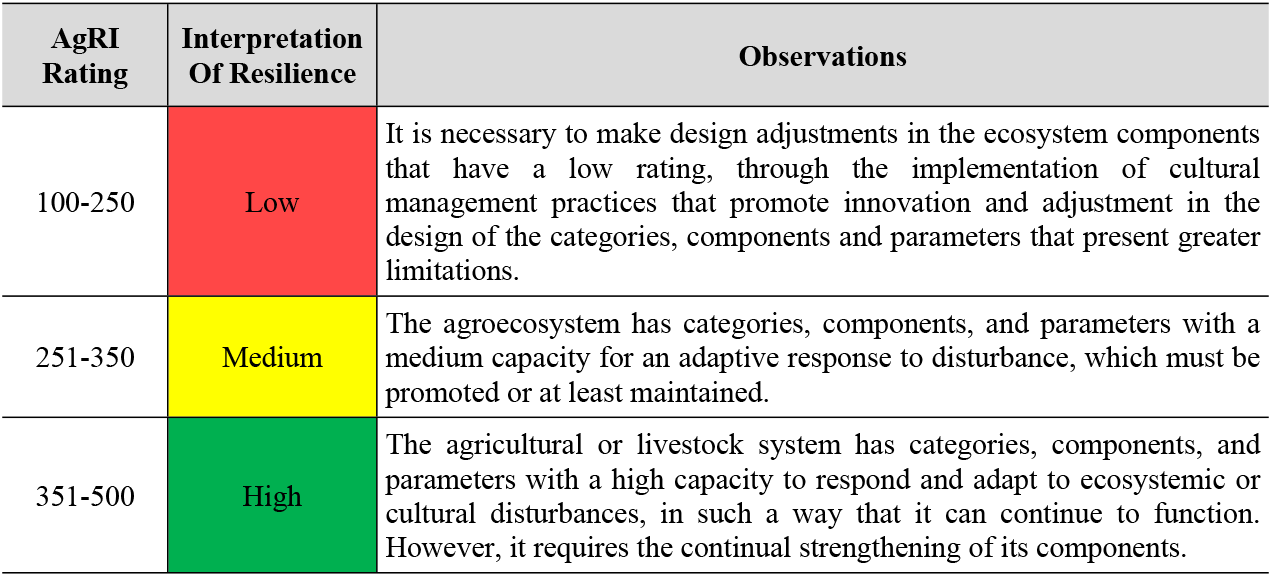
Interpretation of the AgRI (35).

### Results of the application of the AgRI methodology

To verify the relevance, applicability, and robustness of the AgRI, the data related to the typification and classification of citrus production systems in the department of Meta, Colombia (99) were taken into account. The data were obtained in the municipalities of Villavicencio, Granada, Lejanías and Guamal, in the department of Meta, Colombia, where 90% of the planting area and 95% of production is concentrated (**Fig. 5**).

**Figure 5.**
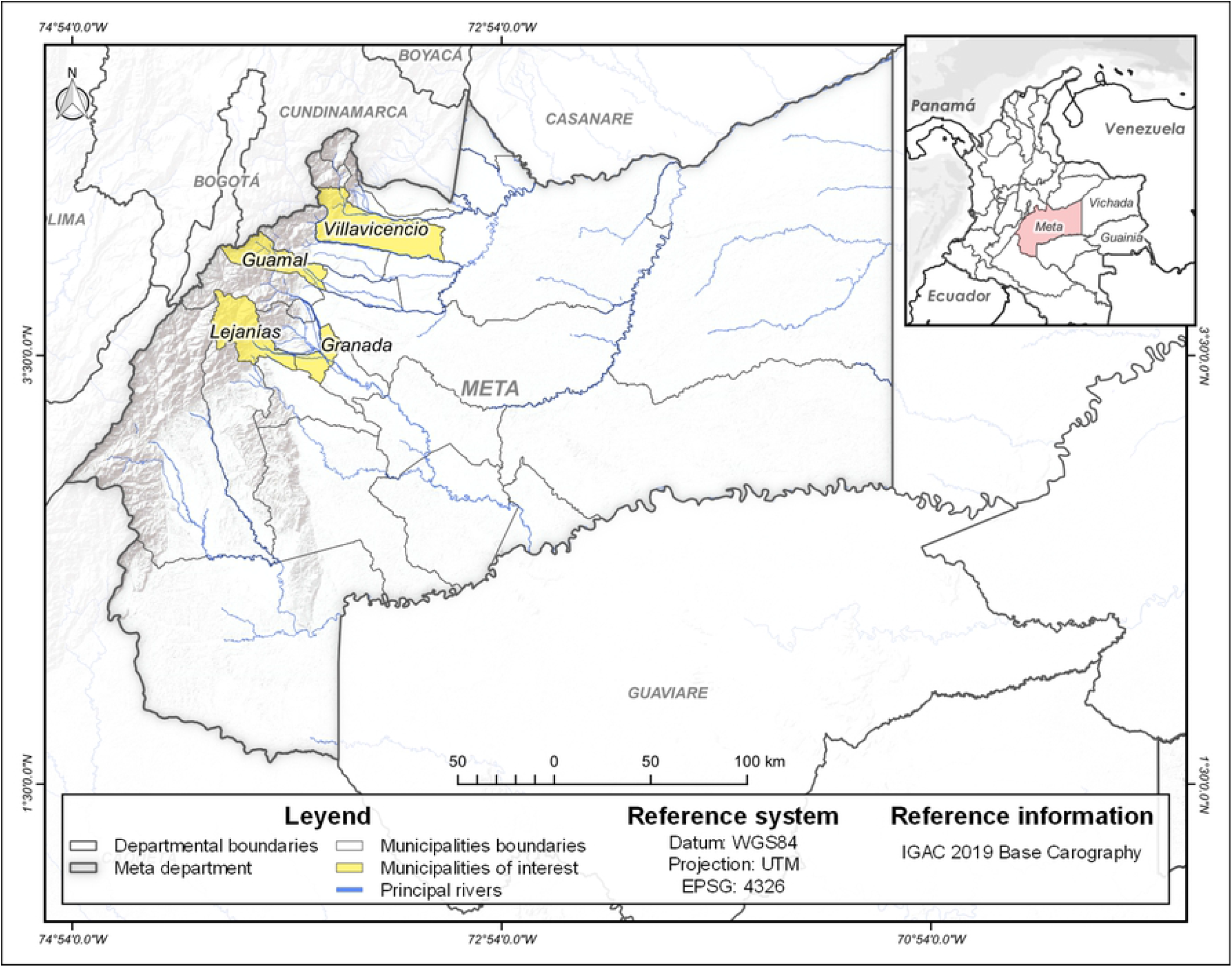
Location of the study area municipalities.

With the collaboration of technicians, farmers, and public and private institutions of a national and regional order, a format (survey) was designed for the compilation of the information to be presented, validated and adjusted in 10 community workshops.

In the selected municipalities, 52 citrus growers were surveyed whose properties had a range of cultural and ecosystem attributes. In total, an area of 710 ha was covered, equivalent to 11.3% of the total planting area.

With the information collected, a database was structured, with which a multivariate statistical analysis was developed. This is considered an ideal technique to simultaneously analyze qualitative and quantitative variables of the production system.

Next, the cluster analysis was carried out based on the Euclidean distance concept using the Ward method. According to this method, the shorter the distance between the productive units, the greater the similarity. Nearby units can therefore belong to the same group while distante units are placed in different groups.

The dendrogram defined the conformation of six (6) groups or “*recommendation domains*” corresponding to groups of farmers, all with attributes of homogeneity within and simultaneously heterogeneity outside of these (**Fig. 6**).

**Figure 6.**
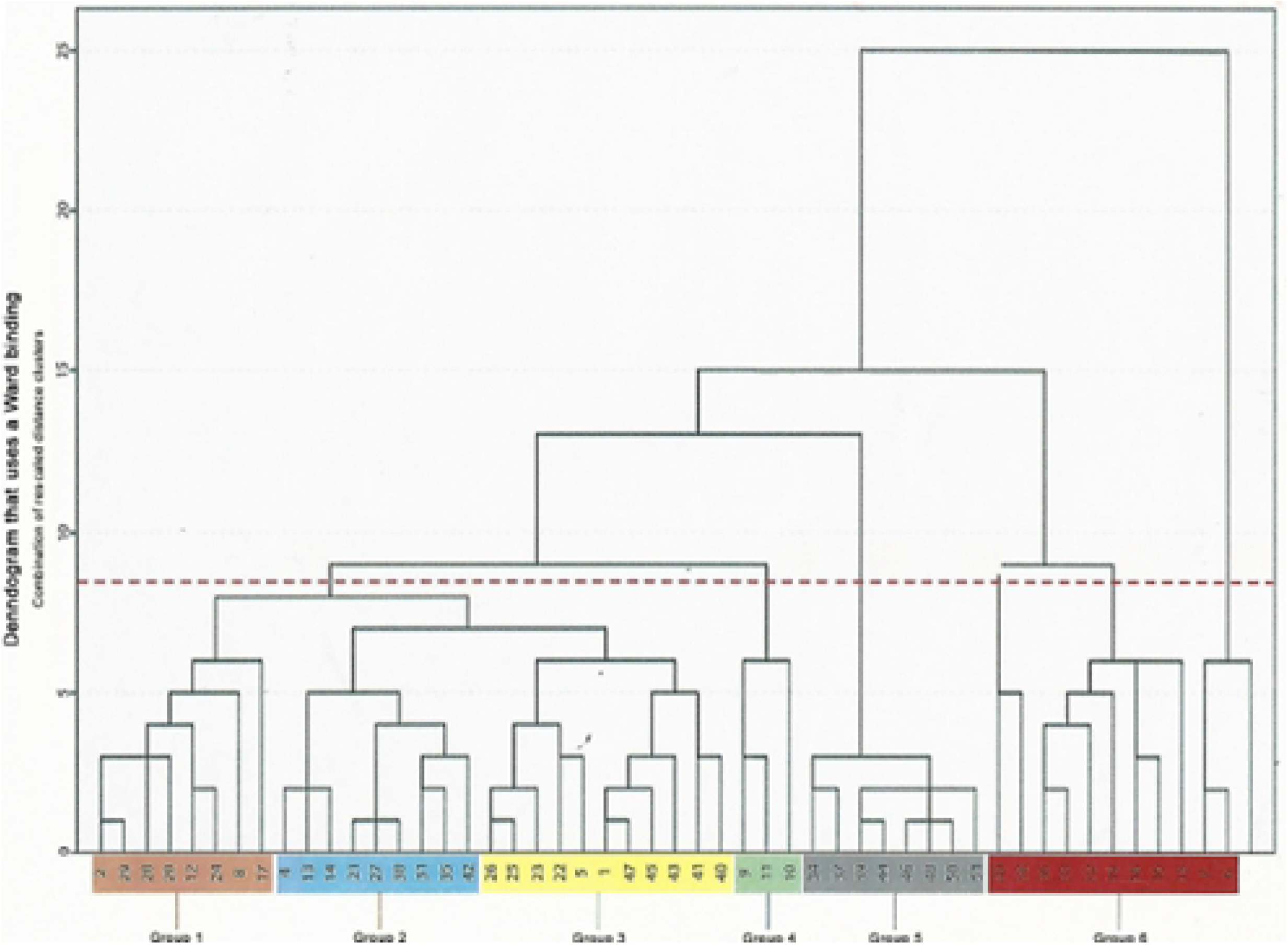
Dendrogram: clustering in six recommendation domains.

In each group or “recommendation domain” of citrus growers, three representative productive units / groups were selected, for a total of 18 farms (n = 18). The respective analyses were carried out from the components of these productive units. The most representative ecosystemic and cultural attributes are listed below in Table 5.

**Table 5.**
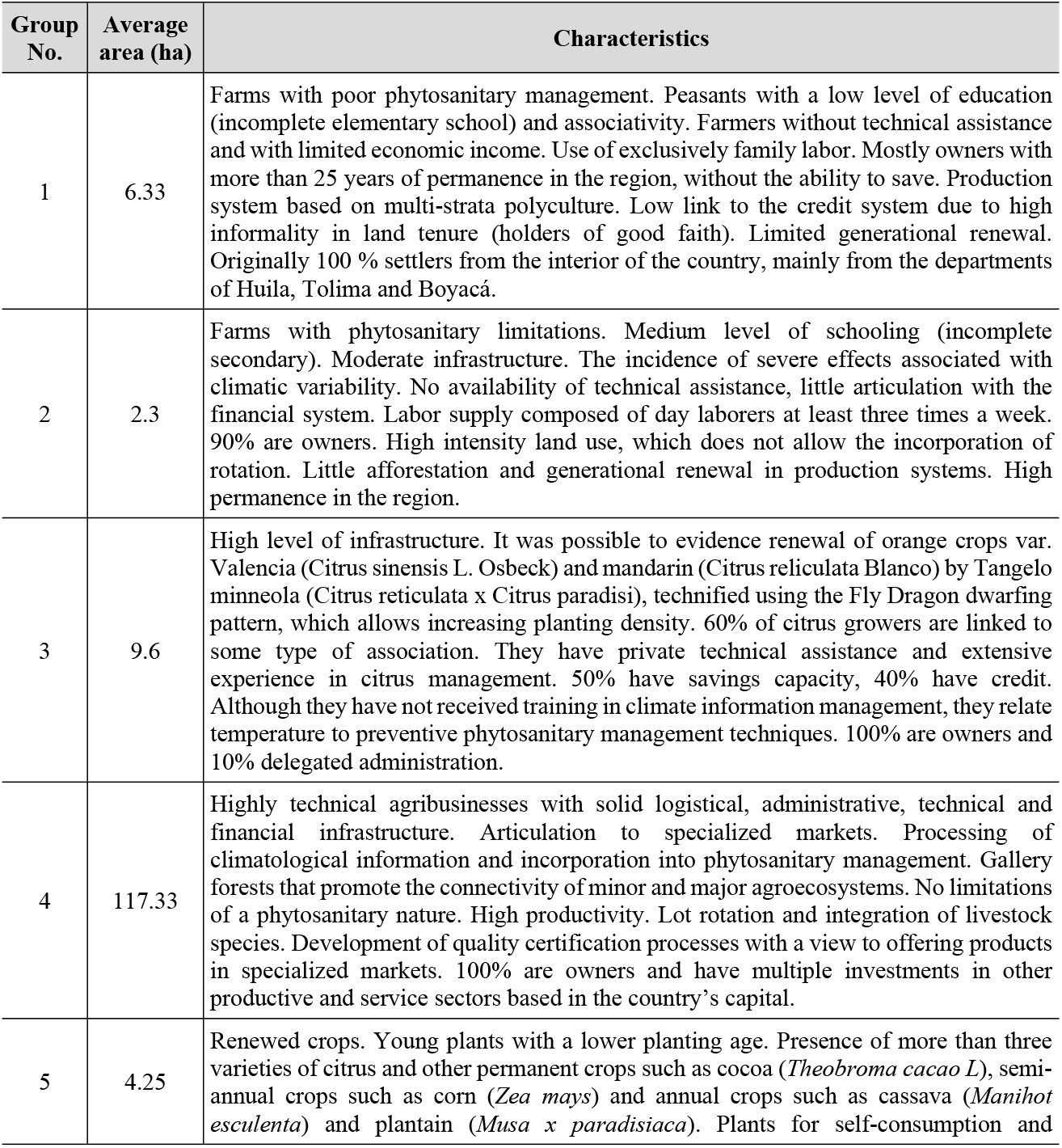
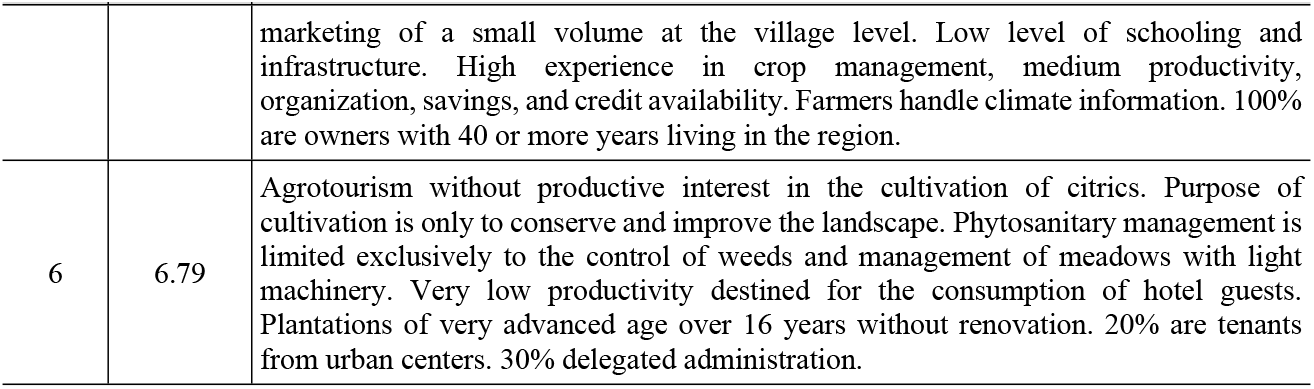
Main attributes of the recommendation domains, in six groups of citrus agroecosystems (18 farms) located in the Department of Meta, Colombia (99)

| Group No. | Average area (ha) | Characteristics |
| --- | --- | --- |
| 1 | 6.33 | Farms with poor phytosanitary management. Peasants with a low level of education (incomplete elementary school) and associativity. Farmers without technical assistance and with limited economic income. Use of exclusively family labor. Mostly owners with more than 25 years of permanence in the region, without the ability to save. Production system based on multi-strata polyculture. Low link to the credit system due to high informality in land tenure (holders of good faith). Limited generational renewal. Originally 100 % settlers from the interior of the country, mainly from the departments of Huila, Tolima and Boyacá. |
| 2 | 2.3 | Farms with phytosanitary limitations. Medium level of schooling (incomplete secondary). Moderate infrastructure. The incidence of severe effects associated with climatic variability. No availability of technical assistance, little articulation with the financial system. Labor supply composed of day laborers at least three times a week. 90% are owners. High intensity land use, which does not allow the incorporation of rotation. Little afforestation and generational renewal in production systems. High permanence in the region. |
| 3 | 9.6 | High level of infrastructure. It was possible to evidence renewal of orange crops var. Valencia (*Citrus sinensis* L. Osbeck) and mandarin (*Citrus reliculata* Blanco) by Tangelo minneola (*Citrus reticulata* x *Citrus paradisi*), technified using the Fly Dragon dwarfing pattern, which allows increasing planting density. 60% of citrus growers are linked to some type of association. They have private technical assistance and extensive experience in citrus management. 50% have savings capacity, 40% have credit. Although they have not received training in climate information management, they relate temperature to preventive phytosanitary management techniques. 100% are owners and 10% delegated administration. |
| 4 | 117.33 | Highly technical agribusinesses with solid logistical, administrative, technical and financial infrastructure. Articulation to specialized markets. Processing of climatological information and incorporation into phytosanitary management. Gallery forests that promote the connectivity of minor and major agroecosystems. No limitations of a phytosanitary nature. High productivity. Lot rotation and integration of livestock species. Development of quality certification processes with a view to offering products in specialized markets. 100% are owners and have multiple investments in other productive and service sectors based in the country's capital. |
| 5 | 4.25 | Renewed crops. Young plants with a lower planting age. Presence of more than three varieties of citrus and other permanent crops such as cocoa (*Theobroma cacao* L), semi-annual crops such as corn (*Zea mays*) and annual crops such as cassava (*Manihot esculenta*) and plantain (*Musa x paradisiaca*). Plants for self-consumption and |
|  |  | marketing of a small volume at the village level. Low level of schooling and infrastructure. High experience in crop management, medium productivity, organization, savings, and credit availability. Farmers handle climate information. 100% are owners with 40 or more years living in the region. |
| 6 | 6.79 | Agrotourism without productive interest in the cultivation of citrics. Purpose of cultivation is only to conserve and improve the landscape. Phytosanitary management is limited exclusively to the control of weeds and management of meadows with light machinery. Very low productivity destined for the consumption of hotel guests. Plantations of very advanced age over 16 years without renovation. 20% are tenants from urban centers. 30% delegated administration. |

### Determination of resilience by group

In the 18 agroecosystems analyzed, the AgRI was calculated for each of the representative farms of the six groups or recommendation domains with the help of (Eq. 4).

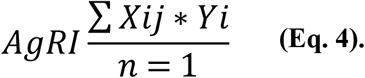

Where:

i= number of evaluated parameters (40).
j= number of farms (18).
X= measurement of the parameter (scale)
y=weighting of the parameter (consultation of experts)

The results are synthesized in Table 6.

**Table 6.** Results of the Agroecosystem Resilience Index (AgRI), in six groups of citrus agroecosystems, located in the Department of Meta, Colombia (35).

| Category | Component | Parameter | Group 1 | Group 2 | Group 3 | Group 4 | Group 5 | Group 6 |
| --- | --- | --- | --- | --- | --- | --- | --- | --- |
| Ecophysiological I | Soil | Slope | 5.61 | 5.61 | 5.61 | 9.35 | 5.61 | 5.61 |
|  |  | Types of erosion | 6.13 | 6.13 | 8.86 | 10.22 | 3.41 | 3.41 |
|  |  | Soil drainage | 6.00 | 6.00 | 7.33 | 10.00 | 3.33 | 3.33 |
|  |  | Effective depth of soil | 6.86 | 5.61 | 8.10 | 9.35 | 4.36 | 3.12 |
|  |  | Fertility management | 6.91 | 6.91 | 8.45 | 6.91 | 6.91 | 5.38 |
|  |  | Landuse LU | 14.74 | 14.74 | 18.01 | 24.57 | 11.46 | 14.74 |
|  |  | Soil conservation practice | 5.22 | 7.30 | 7.30 | 15.65 | 3.13 | 5.22 |
|  |  | Arable Management AM | 1.17 | 1.17 | 3.52 | 5.87 | 3.52 | 1.17 |
|  | Water | Availability of water for irrigation | 9.57 | 5.74 | 21.04 | 28.70 | 17.22 | 9.57 |
|  |  | Water quality and irrigation type | 18.09 | 12.52 | 12.52 | 20.87 | 12.52 | 12.52 |
|  |  | Water conservation practices | 3.17 | 3.17 | 3.17 | 15.87 | 3.17 | 3.17 |
|  | Agrobiodiversity and connectivity | Connection with the Principal Ecological Landscape Structure CPELS | 11.67 | 5.00 | 5.00 | 15.00 | 5.00 | 5.00 |
|  |  | Extension of external connectors EEC | 4.46 | 8.29 | 8.29 | 9.57 | 8.29 | 9.57 |
|  |  | Diversity of external connectors DEC | 15.65 | 13.57 | 7.30 | 15.65 | 7.30 | 11.48 |
|  |  | Extension of internal connectors EIC | 3.26 | 4.57 | 7.17 | 9.78 | 3.26 | 5.87 |
|  |  | Diversity of internal connectors DIC | 11.48 | 9.39 | 11.48 | 15.65 | 7.30 | 11.48 |
| Sociocultural | Capacities | Other management practices OP | 2.87 | 2.87 | 2.87 | 8.61 | 2.87 | 2.87 |
|  |  | Perception – Consciousness (P-C) | 4.86 | 8.74 | 8.74 | 14.57 | 8.74 | 2.91 |
|  |  | Capacity for Action (CA) | 3.00 | 9.00 | 15.00 | 15.00 | 9.00 | 3.00 |
|  | Public services and social security | Potable water | 5.00 | 5.00 | 5.00 | 5.00 | 5.00 | 5.00 |
|  |  | Energy | 5.00 | 5.00 | 5.00 | 5.00 | 5.00 | 5.00 |
|  |  | Health care | 3.67 | 5.00 | 4.33 | 5.00 | 3.00 | 3.00 |
|  | Housing | Quality | 3.00 | 3.00 | 4.33 | 5.00 | 2.33 | 3.67 |
|  | Land structure | Land tenure | 21.52 | 21.52 | 21.52 | 21.52 | 21.52 | 21.52 |
|  |  | Size | 3.65 | 3.65 | 8.52 | 18.26 | 8.52 | 3.65 |
|  | Competencies | Training | 3.12 | 5.61 | 5.61 | 9.35 | 4.36 | 3.12 |
|  |  | Schooling | 1.63 | 2.28 | 1.63 | 4.89 | 2.28 | 2.93 |
|  |  | Organization | 5.12 | 2.20 | 5.12 | 10.98 | 2.20 | 3.66 |
| Economic | Financial capacity | Saving capacity | 5.38 | 5.38 | 8.45 | 11.52 | 3.84 | 11.52 |
|  |  | Productivity | 5.39 | 16.17 | 19.77 | 26.96 | 8.99 | 5.39 |
|  |  | Availability and access to credit | 5.65 | 4.78 | 5.65 | 6.52 | 3.04 | 4.78 |
|  | Market | Destination of production | 8.29 | 11.30 | 11.30 | 11.30 | 9.80 | 2.26 |
|  |  | Trade association | 2.17 | 3.04 | 4.78 | 6.52 | 1.30 | 1.30 |
|  |  | Generation of added value | 5.72 | 10.30 | 10.30 | 10.30 | 10.30 | 3.43 |
| Technological | Practice | Type of cultural practices (management) | 4.78 | 4.78 | 6.70 | 14.35 | 4.78 | 2.87 |
|  |  | Postharvest practices | 6.59 | 2.83 | 8.48 | 10.36 | 8.48 | 2.83 |
|  | Technical assistance | Availability | 1.67 | 1.67 | 4.33 | 5.00 | 1.67 | 1.00 |
|  |  | Type | 1.67 | 1.00 | 3.00 | 5.00 | 1.00 | 1.00 |
|  | Information management | Climatic | 1.96 | 1.96 | 4.57 | 9.78 | 1.96 | 1.96 |
|  |  | Administration | 1.04 | 1.74 | 3.83 | 5.22 | 2.43 | 1.04 |
| Total |  |  | 242.74 | 254.55 | 322.02 | 469.01 | 238.23 | 210.35 |

### Interpretation of the Agroecosystem Resilience Index (AgRI) in six groups of citrus agroecosystems located in the Department of Meta, Colombia

Based on table 4, the results of the Agroecosystemic Resilience Index were interpreted in the 18 farms grouped into six “recommendation domains” (Fig. 7).

**Figure 7.**
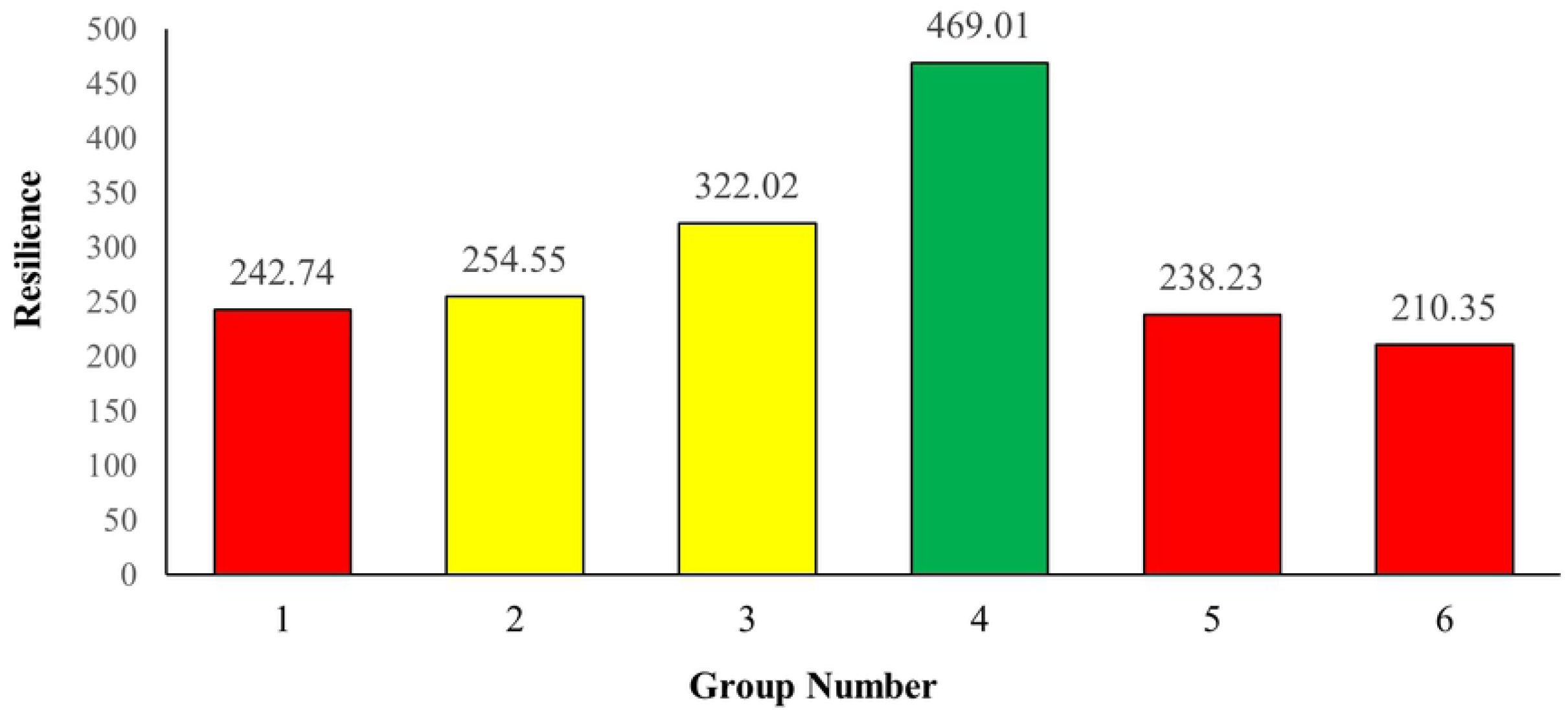
Agroecosystem Resilience Index (AgRI)/group.

According to (35), although ecological systems did not exist in the agroecosystems analyzed, those in group 4 are implementing management practices that are far from the conventional ones, bringing them closer to a typology of agriculture in transition. This group obtained the highest AgRI (469.01). In this case, the parameters that most affected the results were related to the biodiversity of the farms. A wide availability of water resources was manifested in the presence of summer pipes on whose banks gallery forests are developed and conserved, promoting connectivity with the environment (CPELS). There is an integration of larger (bovines and equines) and smaller (beekeeping. poultry) animal species. Cultural readiness is highly significant.

These management conditions, which include the articulation of animal and plant species, promote biodiversity, and represent a good strategy for increasing the resilience of agricultural systems (54,100,101).

This scenario is consistent with the evidence collected by various researchers on the higher levels of resilience of biodiverse agroecosystems compared to conventional ones.

The works of (102,103) stand out as demonstrating that biodiverse agroecosystems adapted and subsequently recovered more quickly and efficiently after the passage of Hurricane Mitch through Central America, with conventional production systems being comparatively more affected.

In contrast, the farms in group 6 present severe limitations in environmental and ecosystem components, according to what is indicated in Tables 5 and 6. In this sense, the result is a low AgRI: 210.35 (red color). The farmers of these farms are more interested in the landscaping service offered by the crops to promote their hotel activities.

The farms in group 3 reached an average AgRI: 322.02 (yellow color), explained in part by their lower availability of water (exclusively deep wells), limited connectivity with their surroundings (CPELS) and few soil conservations practices. The small farm size of these farms’ limits crop rotation practices, due to the intensive use of the soil by farmers. Likewise, they have surpluses that allow them to articulate with the financial system, as well as with the availability of private technical assistance.

The farms of group 2, can be qualitatively classified with an average AgRI of 254.55 (yellow color) as low, being similar to that found in the farms of group 1 and 5 that presented a low AgRI of 242.74 and 238.23 respectively (red color). In this group of farms, significant limitations in the availability and quality of water sources were also observed, in all the parameters of the technological, economic, and sociocultural categories. In this regard, the Colombian state should guide and prioritize programs, plans and incentives so that citrus growers in this group have the possibility of reducing their high vulnerability associated with the low availability of economic resources.

The agricultural systems of group 4 have greater agrobiodiversity, expressed in greater connectivity and diversity of external connectors. They stand out for having greater institutional articulation expressed in different ways: technical assistance, credit support, management of their own marketing channels and generation of added value that allows them to obtain greater resources that they reinvest in technology and ultimately in greater competitiveness.

Cultural and community resilience increases through training processes that strengthen adaptive capacities in farmers, especially in tropical regions where much of the world’s biodiversity is found and must be conserved (104–108).

## CONCLUSIONS

Resilience is an emergent and differential property. The proposal to weight the evaluated variables considers that the ecosystemic and cultural components respond differently to disturbance. Eventually these develop other properties that under normal conditions they would not assume.

The AgRI is an ideal method to assess the resilience of agroecosystems. Its application in the studied systems demonstrated coherence with the characteristics of the environmental components.

It was found that AgRI is associated with agrobiodiversity, which supports the proposed hypothesis that diversified agroecosystems are more resilient.

The AgRI can be used to adjust the design of agroecosystems.

It was found that cultural resilience has a significant impact on the response capacity of agroecosystems to the effect or action of disturbances of a diverse nature. Likewise, it was found that cultural resilience has a significant impact on the response capacity of agroecosystems to the effect or action of disturbances of any nature.

It can be deduced that the implementation of cultural practices executed with a specific purpose fosters functional biodiversity.

The proposed AgRI index is feasible for developing evaluations or analyses oin any agricultural system.

## Acknowledgements

To the Pedagogical and Technological University of Colombia and the project: “Environmental Impact Assessment in Colombia. Critical analysis and Improvement”, Code Hermes: 13129, Universidad Nacional de Colombia, Bogotá and Der Deutsche Akademische Austauschdienst (DAAD) and The Center for Development Research (ZEF), institute of the University of Bonn, Germany. To the citrus producers of the department of Meta, Colombia, for their valuable contributions in the development of the methodology.

## Author contributions

The authors contributed in an equal manner to the research process and the elaboration of the article.

